# Distance-based global analysis of consistent *cis*-bonds in protein backbones

**DOI:** 10.1101/2022.06.11.495244

**Authors:** Tetsuji Okada, Fumiaki Tomoike

**Author notes:** Corresponding author: Tetsuji Okada, Department of Life Science, Gakushuin University, 1-5-1 Mejiro, Toshima-ku, Tokyo 171-8588, Japan.

## Abstract

Biological polypeptides are known to contain *cis*-linkage in their main chain as a minor but important feature. Such anomalous connection of amino acids has different structural and functional effects on proteins. Experimental evidence of *cis*-bonds in proteins is mainly obtained using X-ray crystallography and other methods in the field of structural biology. To date, extensive analyses have been carried out on the experimentally found *cis*-bonds using the Protein Data Bank entry bases and/or residue bases; however, their consistency in each protein has not been examined on a global scale. Data accumulation and advances in methodology enable the use of new approaches from a proteomic point of view. Here, we sought to describe a simple procedure for the detection and confirmation of *cis*-bonds from a set of experimental coordinates for a protein to discriminate this type of bond from isomerizable and/or misassigned bonds. The resulting set of consistent *cis* bonds provides unprecedented insights into the trend of “high *cis* content” proteins and the upper limit of consistent *cis* bonds per polypeptide length. Recognizing such limit would not only be important for a practical check of upcoming structures, but also for the design of novel protein folds beyond the evolutionally-acquired repertoire.

## Introduction

Amino acid connections in proteins, called peptide bonds, mainly assume the *trans* configuration owing to energetical preference. However, deviation from this trend is well-known to occur in a small but meaningful number of cases (Stewart, Sarkar & Wampler, 1990; Craveur et al., 2013). The most prominent evidence of the existence of a *cis*-linkage involves proline in the X-P tandem sequences, where X denotes any amino acids. Non-proline *cis* linkage involves almost any type of two tandem amino acids, including a second *cis*-preferring glycine (Jabs, Weiss & Hilgenfeld, 1999). *Cis*-*trans* bond flip in the main chain could significantly affect the folding manner of a polypeptide. Enzymatically-assisted regulation of this flip has also been evaluated in numerous studies (Göthel & Marahiel, 1999; Fischer & Aumüller, 2003).

Experimentally-determined protein coordinates provide data on the presence of *cis*-bond; however, the correctness of the *cis*/*trans* assignment depends on the resolution of the methods employed. Earlier analyses were performed in a residue-based manner, inevitably due to the limited amount of experimental structure models (Weiss, Jabs & Hilgenfeld, 1998; Pal & Chakrabarti, 1999). However, systematic analysis of *cis* bond inclusion in each protein backbone has not been performed, which might be due to the ability of a substantial fraction of potential *cis* bonds to assume both isomeric configurations according to conditions, such as ligand binding and protein-protein interactions. The presence of hardly isomerizable *cis* bonds in a folded polypeptide, referred to as consistent *cis* bonds in this study, might be advantageous and important for holding the folded polypeptide in a specific conformational state either locally or globally.

We have been collecting backbone distance data for each protein using the so-called distance scoring analysis (DSA) (Anzai et al., 2018), mostly using X-ray crystallographic coordinates archived in Protein Data Bank (PDB) (Burley et al., 2022). This method considers all intramolecular C_α_-C_α_ pair distances for a set of PDB chains per protein in an exactly aligned fashion, and adjacent C_α_-C_α_ distances can be easily extracted from such data. DSA is proposed for quantifying the variability of all intramolecular C_α_-C_α_ pair distances, yielding the score (= average / stdev) for each C_α_-C_α_ pair. Therefore, for a certain peptide bond position in a protein, *cis* only and *cis*/*trans* mixture result in fairly distinct scores, enabling us to infer the consistency of *cis* configuration.

Herein, a large-scale analysis of consistent *cis*-bonds in each protein was performed and confirmed using a redundant set of PDB chains. The results of this analysis were discussed with a particular focus on proteins with high *cis* content per polypeptide and/or per length.

## Methods

### Protein selection

DSA of the PDB coordinate files has been ongoing since 2016 in the order of proteins with larger number of entries. To date, almost all proteins with more than two entries have been processed, with the exception of those residing in large protein complexes and those containing a significant amount of missing C_α_ atoms in their respective full length. A list of the proteins is presented in Table S1 and is included in the top page searchable table of gses2.jp. For most proteins, PDB chains with a single UniProt ID (UniProt Consortium, 2021) are considered a dataset for a protein. Some protein data are derived from chains of more than one UniProt ID, as the sequence identity is more than 95% in the analyzed range. We did not attempt to eliminate the redundancy of a particular type of proteins, such as beta-lactamases in prokaryotic species, HLA antigens in eukaryotic ones, and some proteins in viruses. The average sequence coverage is 82.1% (75.7% and 87.5% for eukaryotic and prokaryotic proteins, respectively) of the full lengths, including signal/pro peptide regions, and the average usage of chains per protein is 11 (13 and 9 for eukaryotic and prokaryotic proteins, respectively).

### *Cis*-bond analysis

DSA was performed as previously described (Anzai et al., 2018; Izumi et al., 2020). A Python script Score-analyzer 3_v05 for Python3 was used to accumulate the basic distance data (dist file) for each protein (Table S2). A sequence range to be analyzed for each protein was reasonably determined to use as many residues and PDB entries as possible, excluding any C_α_ breaks in the range.

In the GUI menu of the Score-analyzer, the *cis*-bond analysis function was implemented. First, a dist file containing all C_α_-C_α_ pair distances was read into the Score analyzer and the scores were calculated (Table S3). The *cis*-bond analysis function identifies a C_α_-C_α_ pair when any one of the pair distances are less than 3.3 Å (likely due to the *cis* configuration), and outputs a list of the pair position number, average distance, and score (Fig. 1, Table S4). Consistent *cis* bonds were selected from the list if the average distance was ∼3.0 Å or less and the score was 20 or more (see below). For a minor fraction of confusing cases, all distance values were individually inspected to determine the validity of the consistency. When some non-bonded (non-tandem position number) pair positions were identified, they were separately counted as remote contacts, and were not considered in the present study. All graphs were generated using Igor Pro (WaveMetrix).

**Fig. 1.**
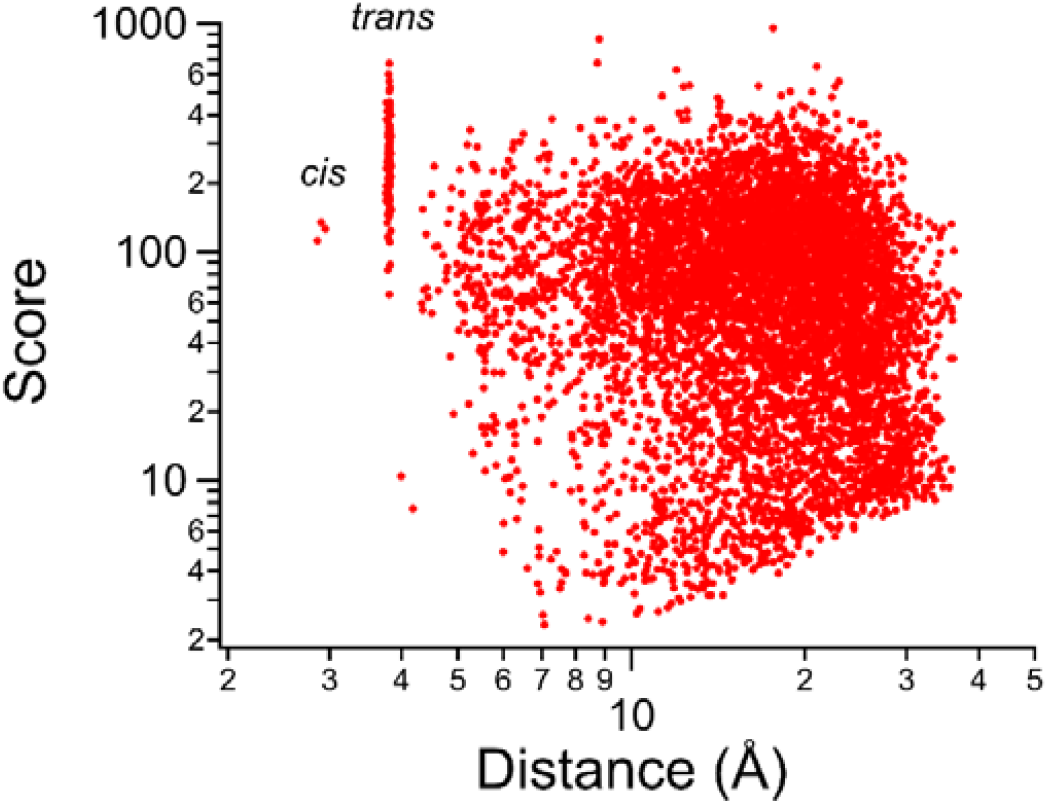
Example log-log main plot of DSA for ribonuclease U2 from *Ustilago sphaerogena*, revealing the presence of three consistent *cis* bonds.

## Results and discussion

### Distance based survey of consistent *cis*-bonds

The main plot and semi-log main plot of a protein obtained from DSA represent the variability of each intramolecular C_α_-C_α_ pair distance (Izumi et al., 2020). As the scores (= average / stdev) were plotted against the average distance, datapoints for adjacent C_α_-C_α_ pairs formed a vertical array at ∼3.8 Å if the peptide bond had the *trans* configuration; however, few datapoints were readily discernible at ∼2.9 Å if the bond was of the *cis* form. Fig. 1 shows an example of the log-log main plot for ribonuclease U2 from *Ustilago sphaerogena* (Smut fungus); a full 114 amino acid chain structure is available from five PDB entries for this protein. Of note, this protein was found to exhibit a high “*cis*/length” feature as described later, with three consistent *cis* bonds (Table S4).

To evaluate the number of peptide bonds with consistent *cis* or mixed *cis*/*trans* configuration in each protein, all C_α_-C_α_ pairs with a distance less than 3.3 Å were detected. The average distance and scores were also listed for the detected pairs as not only the average distance, but also the score, is used. If 10 chains are used for the analysis of a protein, a mixture of one *trans* (3.8 Å) and nine *cis* for a particular bond position results in a score of ∼10 while ten *cis* (distances of 2.85–2.95 Å) results in a score of ∼100. For most cases, scores higher than 20 indicated all *cis*. When a pair with a score ranging from 15 to 20 was observed, inspection of all distance data was performed as the *cis* configuration occasionally exhibits unusually short distance of ∼2.6 Å and such *cis* pair contributes to a substantial lowering of the score to slightly less than ∼20 in some cases. For ribonuclease U2 from *Ustilago sphaerogena*, three consistent *cis* bonds had scores greater than 100 (Fig. 1, Table S4).

### Protein structure set

By the end of February 2023, DSA results were accumulated for 13,422 proteins and a *cis*-bond analysis has been performed for these proteins. Of note, the analyzed proteins are derived from all domains of living organism (eukaryotes, archaea, bacteria, viruses), synthetic constructs and the composition simply reflects the availability of structural coordinates in PDB. As the present DSA only considers continuously-modeled structures, possible effects from bond irregularity due to chain breaks are virtually excluded (Touw, Joosten & Vriend, 2015). However, the protein set is slightly biased toward prokaryotic proteins (7,541 vs 5,129 eukaryotic proteins) owing to fewer chain breaks in the present PDB data. Viral and synthetic proteins account for only 752 of 13,422 proteins (∼5.6%).

All *cis* bond analyses for a protein used at least three chains of identical sequence range and at least two distinct PDB entries in most cases. In contrast to the number of proteins analyzed, the number of used total PDB entries was larger for eukaryotic proteins (54,273) than prokaryotic proteins (43,944), reflecting the higher redundancy of structural studies for the former. The total number of used chains were nearly the same for eukaryotic and prokaryotic proteins (65,559 and 66,682, respectively).

### Consistent *cis* bonds in a protein

The absolute number of *cis*-bonds in a protein might increase as the length of the polypeptide increases. Based on our global analysis, no clear correlation was found (Fig. 2) and the largest number of consistent *cis* bonds for a given polypeptide analyzed to date was 10 in a medium-sized polypeptide, particulate methane monooxygenase subunit B (pmoB), for which 388 continuously modeled amino acids were examined (92.4% of the full length). This protein does not have a UniProt ID but has an NCBI accession of WP_036287217. As a recent cryo-electron microscopy structure (PDB ID: 7S4M) of this subunit only contains six *cis* bonds (Koo et al., 2022), the consistency of these 10 *cis* bonds needs to be confirmed. The second largest number (nine) of consistent *cis* bonds is found in GH43 arabinofuranosidase (AXHd3) of 529 amino acids (97.6% of the full length) from *Humicola insolens*, which also does not have a UniProt ID.

**Fig. 2.**
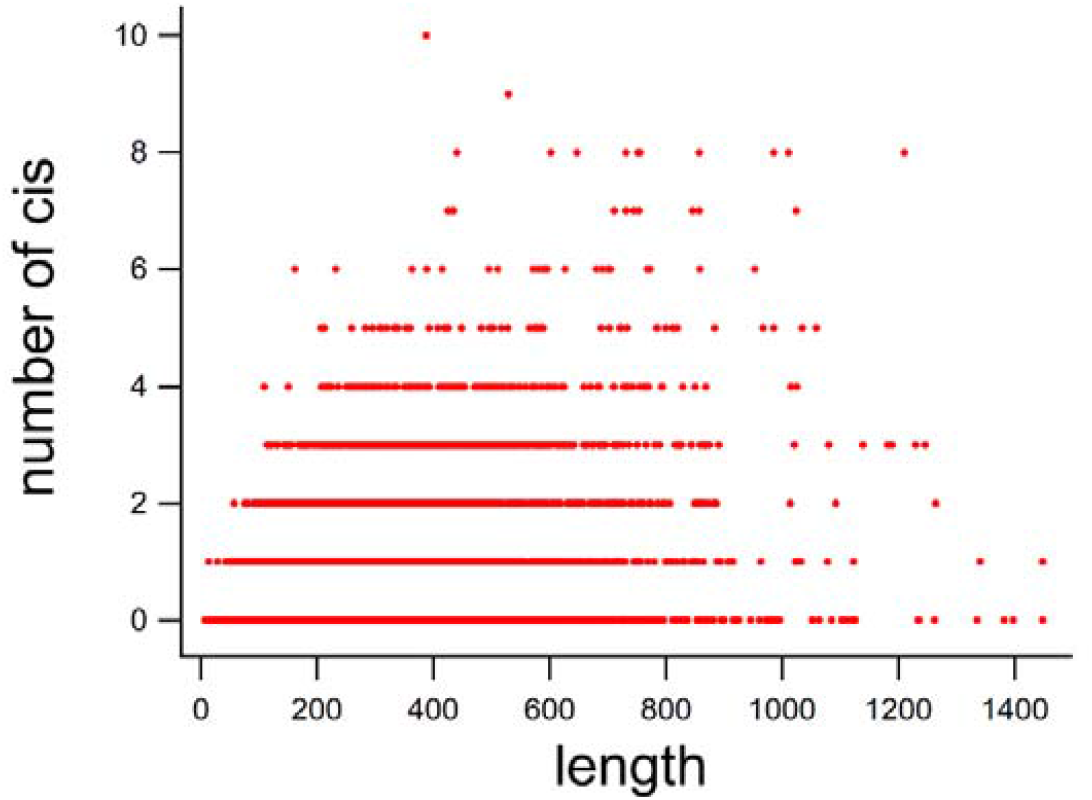
Number of consistent *cis* bonds vs polypeptide length (number of amino acids) for the 13,422 proteins.

Fig. 3a shows the distribution of the number of consistent *cis* bonds in 13,422 proteins, 5,129 eukaryotic proteins, and 7,541 prokaryotic proteins. Not surprisingly, proteins with zero *cis* bonds were most abundant, covering nearly 60% of the total protein. The number of proteins appeared to decrease almost exponentially with increasing *cis* bonds from 1 to 5. Of note, this trend fairly applies to both eukaryotic and prokaryotic proteins. Owing to an insufficient number of proteins containing more than five *cis* bonds, whether this exponential trend extends further is uncertain. Nonetheless, extrapolation of this trend suggests that the probability of finding proteins containing more than 10 consistent *cis* bonds is less than 0.01% (1 out of 10,000), aligning with our finding of only one (pmoB) or no protein with 10 consistent *cis* bonds, and no protein with 11 or more consistent *cis* bonds among the 13,422 proteins. Overall, the average appearance of the consistent *cis* bonds was ∼0.23% (1 per 435 peptide bonds), which is substantially less than that reported in earlier studies (Weiss, Jabs & Hilgenfeld, 1998), in which any consistency in a protein was not examined.

**Fig. 3.**
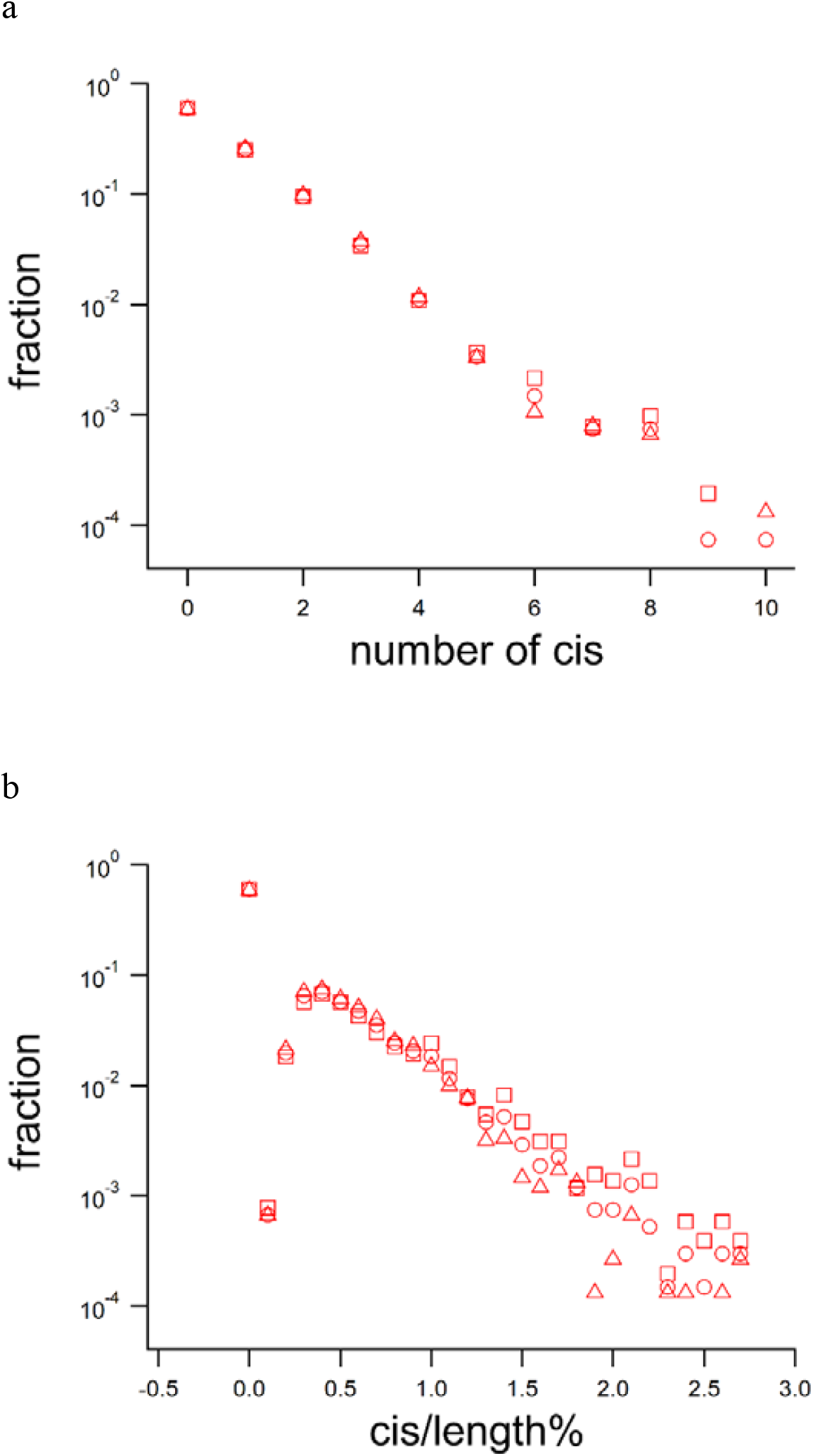
Distribution of protein fraction (○, all proteins; □, eukaryotic proteins; △, prokaryotic proteins). (a) number of consistent *cis* bonds in a continuous modeled polypeptide (b) *cis*/length% for a continuously modeled polypeptide

### High “*cis*/length” protein

As inclusion of *cis* bonds potentially correlates with the folding characteristics of a protein, we analyzed the 13,422 proteins with consistent *cis* percentage (*cis*/length%) calculated by an equation, (number of consistent *cis*) / (amino acid length -1) * 100. Thus, the *cis*/length% becomes 1.0 when one *cis* bond is included in a polypeptide length of 101 amino acids. When the average *cis*/length% for 5,129 eukaryotic and 7,541 prokaryotic proteins was calculated separately, the difference appeared to be marginal (0.25 and 0.22, respectively). Aligning with this finding, the average number of consistent *cis* bonds per protein was similar between eukaryotic and procaryotic proteins (0.63 and 0.65, respectively). Protein distribution of the *cis*/length% is shown in Fig. 3b, demonstrating a peak around 0.5%.

Fig. 4 shows the plot of *cis*/length% against length for all, eukaryotic, and prokaryotic proteins. Each dot (data of a protein) forming a curved line belongs to a group of the same number of consistent *cis* bonds (flat bottom dots represent zero *cis* bond proteins) in the figure. The pattern of all dots is consistent with the above findings; the present upper limit of the number of consistent *cis* bonds per protein is 10, and near exponential decay of protein fraction with increasing number of consistent *cis* bonds. Of note, most of the examined proteins had a *cis*/length% of less than 2.0.

**Fig. 4.**
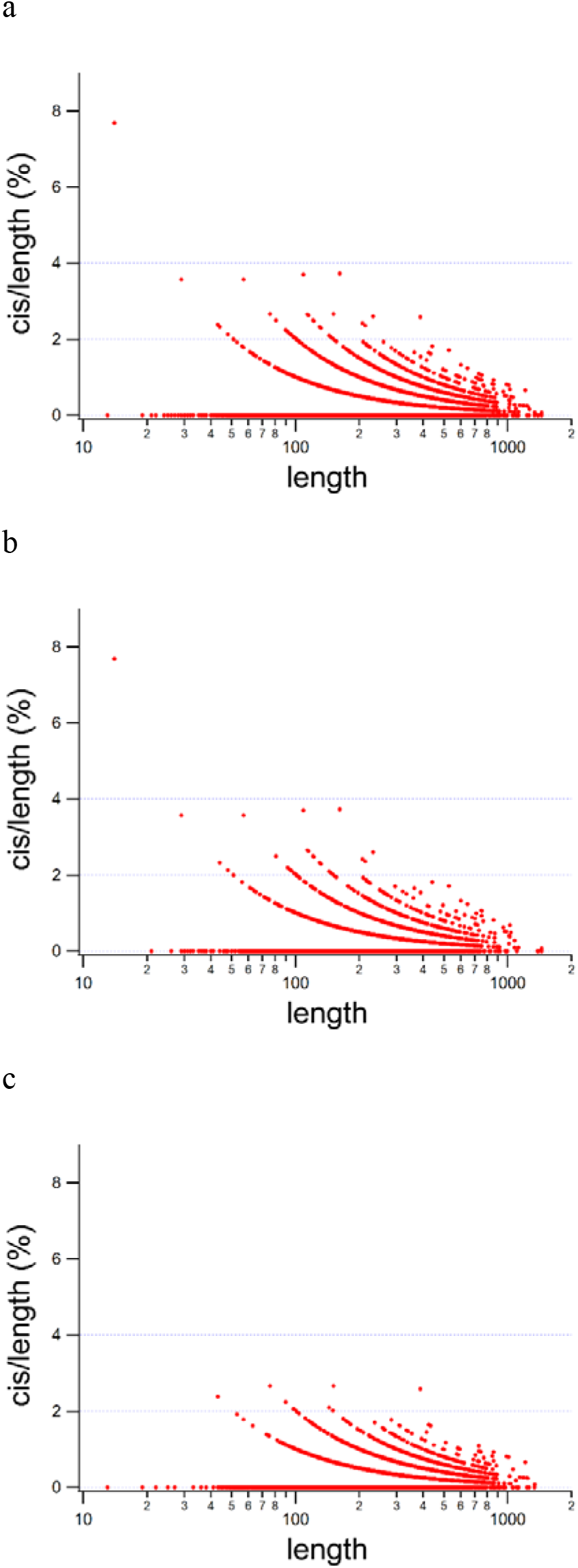
Polypeptide length dependency of the number of consistent *cis* bonds in continuously modeled proteins. (a) all proteins (b) eukaryotic proteins (c) prokaryotic proteins

Of the 13,422 proteins, only 39 were found to have a *cis*/length% of more than 2.0; these proteins are listed in Table 1 as high *cis*/length% proteins. Interestingly, 30 of the 39 high *cis*/length% proteins are of eukaryotic origin. The most abundant legacy Pfam family in eukaryotic high *cis*/length% proteins is V-set (5 of 30) under the clan of Ig. As there is an additional protein under Ig clan in this list, six immunoglobulin fold proteins significantly contribute to this eukaryotic high *cis*/length% proteins. However, the *cis*/length% of these Ig clan proteins do not exceed 3.0 in the present analysis.

**Table 1.** High *cis*/length% proteins

| Gene | Protein | UniProt | Length | Length(%) | *cis*/length% |
| --- | --- | --- | --- | --- | --- |
| SFTI1 | Trypsin inhibitor 1 | Q4GWU5 | 14 | 100.0 | 7.69231 |
| GM2A | Ganglioside GM2 activator | P17900 | 162 | 83.9 | 3.72671 |
| n.a. | Mannose/sialic acid-binding lectin | Q8L568 | 109 | 68.1 | 3.70370 |
| OAK1 | Kalata-B1 | P56254 | 29 | 100.0 | 3.57143 |
| n.a. | Bowman-Birk type proteinase inhibitor | P01055 | 57 | 80.3 | 3.57143 |
| TUSA | Sulfur carrier protein TusA | P0A892 | 76 | 93.8 | 2.66667 |
| yojM | Superoxide dismutase-like protein YojM | O31851 | 151 | 77.0 | 2.66667 |
| RNU2 | Ribonuclease U2 | P00654 | 114 | 100.0 | 2.65487 |
| Ctla4 | Cytotoxic T-lymphocyte protein 4, extracellular | P09793 | 115 | 51.6 | 2.63158 |
| KIR3DL1 | Killer cell immunoglobulin-like receptor 3DL1, extracellular | P43629 | 232 | 52.3 | 2.59740 |
| n.a. | Particulate Methane monooxygenase subunit B (pmoB) | n.a. | 388 | 92.4 | 2.58398 |
| Vapb | Vesicle-associated membrane protein-associated protein B | Q9QY76/O95292 | 121 | 49.8 | 2.50000 |
| n.a. | 6.5 kDa glycine-rich antifreeze protein | Q38PT6 | 81 | 100.0 | 2.50000 |
| n.a. | Major sperm protein isoform alpha | P27439 | 122 | 96.1 | 2.47934 |
| crchiA | Chitinase A | Q4W6L6 | 207 | 53.5 | 2.42718 |
| RPO12 | DNA-directed RNA polymerase subunit P | B8YB64 | 43 | 89.6 | 2.38095 |
| Cdh2 | Cadherin-2 | P15116 | 213 | 23.5 | 2.35849 |
| NPC2 | NPC intracellular cholesterol transporter 2 | P79345 | 130 | 100.0 | 2.32558 |
| n.a. | Monellin chain A | P02881 | 44 | 97.8 | 2.32558 |
| ret | Ribonuclease mitogillin | P67876 | 132 | 88.6 | 2.29008 |
| TTHB138 | FeS_assembly_P domain-containing protein | Q53W28 | 90 | 87.4 | 2.24719 |
| Trav11 | T cell receptor alpha variable 11D | A0A0B4J1J9 | 92 | 98.9 | 2.19780 |
| SLL-2B | Galactose-binding lectin | A4CYJ6 | 94 | 79.0 | 2.15054 |
| n.a. | Monellin chain B | P02882 | 48 | 96.0 | 2.12766 |
| IGKV1D-33 | Immunoglobulin kappa variable 1D-33 | P01593 | 95 | 100.0 | 2.12766 |
| GM10881 | Ig kappa chain V-V region L7 (Fragment) | P01642 | 95 | 100.0 | 2.12766 |
| Lair1 | Leukocyte-associated immunoglobulin-like receptor 1, extracell | Q8BG84 | 96 | 36.5 | 2.10526 |
| IGKV3-20 | Immunoglobulin kappa variable 3-20 | P01619 | 96 | 100.0 | 2.10526 |
| LOC109719601 | Jacalin-like lectin | Q53J09 | 144 | 99.3 | 2.09790 |
| fabZ | 3-hydroxyacyl-[acyl-carrier-protein] dehydratase FabZ | A1KRL1 | 144 | 96.6 | 2.09790 |
| n.a. | BEL-beta trefoil | R4GRU5 | 146 | 100.0 | 2.06897 |
| n.a. | Heat-labile enterotoxin IIB, B chain | P43529 | 98 | 99.0 | 2.06186 |
| LT-IIc1 B | Heat-labile enterotoxin IIA, B chain | H6W8F2 | 98 | 100.0 | 2.06186 |
| PETE | Plastocyanin, chloroplastic | P07030 | 99 | 100.0 | 2.04082 |
| PETE | Plastocyanin A, chloroplastic | P00299 | 99 | 100.0 | 2.04082 |
| PETE_2 | Plastocyanin B, chloroplastic | P11970 | 99 | 100.0 | 2.04082 |
| PETE | Plastocyanin, chloroplastic | P00289 | 99 | 100.0 | 2.04082 |
| n.a. | Frutapin | A0A2D0TC52 | 150 | 100.0 | 2.01342 |
| RUS | Rusticyanin | P0C918 | 150 | 96.8 | 2.01342 |

Numerals with red color in the length(%) column are the values calculated without signal/pro peptide region at the respective N-terminal.

Only five proteins have a *cis*/length% higher than 3.0 (Fig. 4, Table 1) and these proteins are from eukaryotic species (one human and four plant proteins). Based on the dot distribution in Fig. 4, these proteins appear to be rather exceptional. In fact, two of these proteins (SFTI1 and OAK1) form a cyclic polypeptide structure. SFTI1 and Non cyclic Bowman-Birk type proteinase inhibitor (P01055) are classified in the same Pfam family of Bowman-Birk_leg, and P01055 is extensively disulfide-bonded (seven S-S bonds in 57 amino acid chain). Whereas the consistency analysis of P01055 protein was carried out for 57 amino acids (80.3% of full length without signal/pro peptide), the longest model structure (PDB ID: 5J4Q) suggests that the cis/length% would be ∼2.857 (2 cis in 71 residues). GM2A, with a novel β-cup topology, contains six consistent *cis* bonds in 162 amino acid chain (Wright, Li & Rastinejad, 2000) and belongs to the E1_DerP2_DerF2 Pfam family. Both E1_DerP2_DerF2 and Bowman-Birk_leg families were mainly found in eukaryotes. Six *cis* bonds were also found in the single PDB entry (2AGC) for the mouse homolog (75% sequence identity) of this protein, and both human and mouse GM2A contains four disulfide bonds. Mannose/sialic acid-binding lectin (Q8L568) from *Polygonatum cyrtonema* does not contain any Pfam domains, but belongs to the *Galanthus nivalis* agglutinin (GNA)-related lectin family with beta-prism II fold (Ding et al., 2010). As the fourth *cis* bond of this protein is located at the edge of the truncated C-terminal, the number of consistent *cis* might be three (*cis*/length% is ∼2.778). Therefore, we suppose at this point that GM2A is the only non-cyclic protein with a *cis*/length% higher than 3.0. The proline content per length of GM2A was fairly high (∼9.88%) compared with the reported average (Morgan & Rubenstein, 2013), but not outstandingly high compared to that of other proteins in Table 1.

### Consistent *cis* bonds with proline and others

X-P is well known as the most frequently found amino acid pair that forms the *cis* configuration, where X stands for any amino acids and P stands for proline. All consistent *cis* bonds in the top five *cis*/length% proteins described above are of the X-P type. Only five of the top 39 *cis*/length% proteins listed in Table 1 contain non-X-P type *cis* bonds that are known to occur very rarely (Richardson et al., 2018). All high *cis*/length% vertebrate proteins (11 of 39) only contain X-P *cis* bonds. pmoB protein with 10 tentative consistent *cis* bonds in X-ray structures (6 in cryo-EM) contains seven non-XP types, and the remaining three X-P *cis* bonds are found in the cryo-EM structure.

For high *cis* number proteins, such as those containing more than five consistent *cis* bonds, approximately one-third of the total *cis* bonds is of the non X-P type (96 of 289). This apparent bias is partly due to the abundance of beta-galactosidases from various species in high *cis* number proteins.

### Mixed *cis*/*trans* bonds

Based on our analysis, 6,921 sites were assigned as mixed *cis*/*trans* configuration from 13,422 proteins. This amount may not be trivial as it is only ∼18.6% less than 8,499 of consistent *cis* sites from the same protein set. One of the likely reasons for this finding is the lack of a resolution cut off. Resolution dependent misassignment could occur in either directions, *trans* to *cis* or *cis* to *trans*, as revealed previously (Weiss, Jabs & Hilgenfeld, 1998; Lorenzen et al., 2005; Croll, 2015).

The mixed sites were roughly classified into three categories: *cis*-dominant, *trans*-dominant, and evenly mixed, as according to the average distances of closer to 2.9 Å, 3.8 Å, and 3.3 Å, respectively. As a site of mixed *cis* bond was identified if one or more of its distances of less than 3.3 Å were contained in a set of chains used for a protein, a significant amount of *trans*-dominant bonds was found to reside at 6,921 bond sites. Many of these bonds are the results of misassignment as *cis* in one or few chains. On the other hand, some of the *cis*-dominant bonds might be better assigned as consistent *cis* as misassignment to *trans* could also occur. Nonetheless, such cases, even if present, do not affect the main findings of this study. In fact, inspection of all distance data for confusing cases indicated that only ∼0.45% (∼60 in 13,422) of proteins might contain more consistent *cis* bonds than that found in the present study.

Finally, evenly mixed bonds might reflect the convertible nature of the sites depending on the conditions, such as ligand-binding and protein-protein interaction (Joseph, Srinivasan & de Brevern, 2012). Further detailed analysis on the mixed sites should be performed in the future.

## Conclusions

In the present analysis, we analyzed 104,854 PDB entries and 141,017 chains from these entries, and showed that the percentage of consistent *cis* bonds in a protein would rarely exceed three. For a more complete analysis on the PDB data archive, single entry proteins comprising three or more chains must be evaluated. An analysis of two entry proteins with only three or four chains is in progress. In addition, multiprotein complex entries have not been extensively used. In many cases, only one or a few proteins (with larger coverage of the full length) in a complex are included in the present analysis. Furthermore, many proteins remain unanalyzed due to the presence of extensive chain breaks and/or poor sequence coverage in the available coordinate files.

The level of confidence for the consistent *cis* bonds in the present analysis inevitably varies for the 13,422 proteins as the number of usable chains is limited by their availability from the PDB archive. The most confident *cis* bonds may be two positions (29–30 and 200–201) in human carbonic anhydrase 2 determined from 1,015 chains and 1,015 entries. The present analysis also revealed four mixed sites (28–29, 233–234, 234–235, and 236–237) for this protein, all of which were *trans* dominant. On the other hand, the recently reported carbonic anhydrase (A0A3Q0KSG2) from *Schistosoma mansoni* (Angeli et al., 2021), in which four consistent *cis* bonds and no mixed sites were found, was analyzed using 8 entries and 13 chains. Thus, the confidence of these four *cis* bonds might be considered less than that of the human homolog.

Of note, once we obtained the number of confident cis bonds by the present method, future re-analysis by adding new experimental coordinates would not increase it. It is only possible that any of these consistent *cis* bonds could be re-assigned as mixed sites depending on the added coordinates. Therefore, the upper limit of *cis*/length% value is already very reliable for the proteins in which no mixed sites were found. Recognizing such limit would not only be important for a practical check of upcoming structures, but also for the design of novel protein folds beyond the evolutionally-acquired repertoire.

## References

Angeli A, Ferraroni M, Da’dara AA, Selleri S, Pinteala M, Carta F, Skelly PJ, Supuran CT. 2021. Structural Insights into Schistosoma mansoni Carbonic Anhydrase (SmCA) Inhibition by Selenoureido-Substituted Benzenesulfonamides. Journal of Medicinal Chemistry 64:10418–10428. DOI: 10.1021/acs.jmedchem.1c00840.

Anzai R, Asami Y, Inoue W, Ueno H, Yamada K, Okada T. 2018. Evaluation of variability in high-resolution protein structures by global distance scoring. Heliyon 4:e00510. DOI: 10.1016/j.heliyon.2018.e00510.

Burley SK, Bhikadiya C, Bi C, Bittrich S, Chen L, Crichlow GV, Duarte JM, Dutta S, Fayazi M, Feng Z, Flatt JW, Ganesan SJ, Goodsell DS, Ghosh S, Kramer Green R, Guranovic V, Henry J, Hudson BP, Lawson CL, Liang Y, Zardecki C. 2022. RCSB Protein Data Bank: Celebrating 50 □ years of the PDB with new tools for understanding and visualizing biological macromolecules in 3D. Protein Science 31:187–208. DOI: 10.1002/pro.4213.

Craveur P, Joseph AP, Poulain P, de Brevern AG, Rebehmed J. 2013. Cis-trans isomerization of omega dihedrals in proteins. Amino Acids 45:279–289. DOI: 10.1007/s00726-013-1511-3.

Croll TI. 2015. The rate of cis-trans conformation errors is increasing in low-resolution crystal structures. Acta Crystallographica. Sect. D, Biological Crystallography 71:706–709. DOI: 10.1107/S1399004715000826.

Ding J, Bao J, Zhu D, Zhang Y, Wang D-C. 2010. Crystal structures of a novel anti-HIV mannose-binding lectin from Polygonatum cyrtonema Hua with unique ligand-binding property and super-structure. Journal of Structural Biology 171:309–317. DOI: 10.1016/j.jsb.2010.05.009.

Fischer G, Aumüller T. 2003. Regulation of peptide bond cis/trans isomerization by enzyme catalysis and its implication in physiological processes. Reviews of physiology, biochemistry and pharmacology 148:105–150. DOI: 10.1007/s10254-003-0011-3.

Göthel SF, Marahiel MA. 1999. Peptidyl-prolyl cis-trans isomerases, a superfamily of ubiquitous folding catalysts. Cellular and Molecular Life Sciences 55:423–436. DOI: 10.1007/s000180050299.

Izumi K, Saho E, Kutomi A, Tomoike F, Okada T. 2020. Repertoire of morphable proteins in an organism. PeerJ 8:e8606. DOI: 10.7717/peerj.8606.

Jabs A, Weiss MS, Hilgenfeld R. 1999. Non-proline cis peptide bonds in proteins. Journal of Molecular Biology 286:291–304. DOI: 10.1006/jmbi.1998.2459.

Joseph AP, Srinivasan N, de Brevern AG. 2012. Cis-trans peptide variations in structurally similar proteins. Amino Acids 43:1369–1381. DOI: 10.1007/s00726-011-1211-9.

Koo CW, Tucci FJ, He Y, Rosenzweig AC. 2022. Recovery of particulate methane monooxygenase structure and activity in a lipid bilayer. Science 375:1287–1291. DOI: 10.1126/science.abm3282.

Lorenzen S, Peters B, Goede A, Preissner R, Frömmel C. 2005. Conservation of cis prolyl bonds in proteins during evolution. Proteins 58:589–595. DOI: 10.1002/prot.20342.

Morgan AA, Rubenstein E. 2013. Proline: the distribution, frequency, positioning, and common functional roles of proline and polyproline sequences in the human proteome. Plos One 8:e53785. DOI: 10.1371/journal.pone.0053785.

Pal D, Chakrabarti P. 1999. Cis peptide bonds in proteins: residues involved, their conformations, interactions and locations. Journal of Molecular Biology 294:271–288. DOI: 10.1006/jmbi.1999.3217.

Richardson JS, Williams CJ, Hintze BJ, Chen VB, Prisant MG, Videau LL, Richardson DC. 2018. Model validation: local diagnosis, correction and when to quit. Acta crystallographica. Section D, Structural biology 74:132–142. DOI: 10.1107/S2059798317009834.

Stewart DE, Sarkar A, Wampler JE. 1990. Occurrence and role of cis peptide bonds in protein structures. Journal of Molecular Biology 214:253–260. DOI: 10.1016/0022-2836(90)90159-J.

Touw WG, Joosten RP, Vriend G. 2015. Detection of trans-cis flips and peptide-plane flips in protein structures. Acta Crystallographica. Sect. D, Biological Crystallography 71:1604–1614. DOI: 10.1107/S1399004715008263.

UniProt Consortium. 2021. UniProt: the universal protein knowledgebase in 2021. Nucleic Acids Research 49:D480–D489. DOI: 10.1093/nar/gkaa1100.

Weiss MS, Jabs A, Hilgenfeld R. 1998. Peptide bonds revisited. Nature Structural Biology 5:676. DOI: 10.1038/1368.

Wright CS, Li SC, Rastinejad F. 2000. Crystal structure of human GM2-activator protein with a novel beta-cup topology. Journal of Molecular Biology 304:411–422. DOI: 10.1006/jmbi.2000.4225.

